# Humans rely more on talker identity than temporal coherence in an audiovisual selective attention task using speech-like stimuli

**DOI:** 10.1101/2022.08.18.503976

**Authors:** Madeline S Cappelloni, Vincent S Mateo, Ross K Maddox

## Abstract

Audiovisual integration of speech can benefit the listener by not only improving comprehension of what a talker is saying but also helping a listener pick a particular talker’s voice out of a mix of sounds. Binding, an early integration of auditory and visual streams that helps an observer allocate attention to a combined audiovisual object, is likely involved in audiovisual speech processing. Although temporal coherence of stimulus features across sensory modalities has been implicated as an important cue for non-speech stimuli (Maddox et al., 2015), the specific cues that drive binding in speech are not fully understood due to the challenges of studying binding in natural stimuli. Here we used speech-like artificial stimuli that allowed us to isolate three potential contributors to binding: temporal coherence (are the face and the voice changing synchronously?), articulatory correspondence (do visual faces represent the correct phones?), and talker congruence (do the face and voice come from the same person?). In a trio of experiments, we examined the relative contributions of each of these cues. Normal hearing listeners performed a dual detection task in which they were instructed to respond to events in a target auditory stream and a visual stream while ignoring events in a distractor auditory stream. We found that viewing the face of a talker who matched the attended voice (i.e., talker congruence) offered a performance benefit. Importantly, we found no effect of temporal coherence on performance in this task, a result that prompts an important recontextualization of previous findings.

## Introduction

When we’re struggling to understand a talker in a noisy environment, one of the most helpful strategies is to look at the talker’s face. Numerous studies have touted the benefits of faces to speech comprehension (Arnold & Hill, 2001; Reisberg et al., 1987; Sumby & Pollack, 1954). Yet, in principle, a face that matches the talker’s voice could do more than improve our comprehension of their words, it might also improve our ability to attend that voice when it’s presented in a mix of sounds.

### Binding supports object-based attention

Audiovisual binding is the mechanism by which stimulus features are combined across modalities into a single object that can be attended by the observer (Bizley et al., 2016). It is thought to be an automatic and early computation that occurs independently of an observer’s goals. Importantly, binding, sometimes called “early integration,” is a separate mechanism from what is traditionally conceived of as late integration. Late integration is often modeled as ideal Bayesian causal inference, in which unisensory estimates of a stimulus are combined based on their relative reliabilities if the observer infers that they likely arise from a common cause (Körding et al., 2007).

Late integration allows the observer to improve the accuracy of their perceptual judgments by combining redundant estimates across sensory modalities. However, binding is thought to benefit an observer by allowing them to focus attention on a combined audiovisual object, rather than dividing attention between separate auditory and visual objects. Importantly, forming an audiovisual object allows the observer to not only attend the features that correspond across modalities (the perception of which is also improved by late integration), but also those that are only represented in one modality (orthogonal cues). Tests for audiovisual binding then exploit this key difference between audiovisual binding and late integration—that only audiovisual binding should impact the perception of orthogonal cues (Bizley et al., 2016; Maddox et al., 2015).

### Temporal coherence has been shown to drive binding

Previously, Maddox et al. (2015) designed a task in which subjects were instructed to detect pitch or timbre changes in a target auditory stream (a tone complex) while ignoring a masker (a second tone complex) and also responding to color changes in a visual stream (a circle). When the size of the visual shape and amplitude of the target auditory stream were comodulated with the same temporal envelope, listeners improved their sensitivity to orthogonal events in the auditory stream, suggesting that temporal coherence was a potent cue for binding (Maddox et al., 2015). This result was later replicated in Atilgan and Bizley (2020).

In Fiscella et al. (2022), we aimed to show that binding occurred not only for circles and tone complexes, but also had a role in audiovisual speech perception. In this experiment, we asked subjects to discriminate whether the target sentence had a pitch modulation or not while ignoring a masker sentence and coincidentally performing a visual catch task. We considered four visual conditions. The speech video could match the target sentence, and thus be temporally coherent, or could match the masker sentence, making them temporally incoherent. The videos were also presented upright or inverted, where the inverted videos had disrupted visual articulatory cues (i.e., the canonical mouth shapes associated with different speech sounds were upside down). We found that video matching the target sentence was beneficial, but the orientation of the video did not matter (even though in an earlier part of that study we had demonstrated an intelligibility in noise deficit arising from the inversion). These results suggested that temporal coherence, but not articulatory correspondence contributed to binding. However, there were several differences between the original task in Maddox et al (2015) and this study due to restrictions imposed by the natural speech stimuli, leading to lingering curiosity about cues that can influence binding in speech and ultimately motivating the current study.

### The implications of binding’s neural underpinnings

Even prior to work investigating audio-visual binding, studies have investigated evidence that audio-visual integration could occur as early as primary auditory cortex (A1) (Schroeder et al., 2008), an area that is traditionally considered to be unisensory. A variety of imaging modalities in humans have corroborated this theory. fMRI has shown activation of A1 to visual speech (Pekkola et al., 2005). EEG studies have shown that audio-visual speech influences ERPs at latencies consistent with A1 (Besle et al., 2004; van Wassenhove et al., 2005). MEG work also has implicated A1 specifically in the enhancement of speech envelope when viewing a matching face (Golumbic et al., 2013).

Atilgan et al. (2018) built on this work by recording directly from A1 in ferrets. They showed that activity in A1 in response to stimuli from Maddox et al. (2015) depends on which visual stimulus is presented with the sounds. When presented with a mix of sounds and a visual stimulus that was coherent with one of the sounds, activity in A1 preferentially represented that sound. Not only did this work strengthen the evidence that temporal coherence is an important cue for binding, but also suggested that binding occurs in areas that have access primarily to low level cues.

### Audiovisual correspondences in speech

Here we considered three separate audiovisual correspondences that could drive binding in speech: temporal coherence, articulatory congruence, and talker congruence. The first, temporal coherence, is the low-level physical relationship between the timecourses of mouth movements and sound production. In natural speech the visual area of the mouth is roughly correlated with the amplitude of the sound, specifically of energy in a frequency range corresponding with the second formant (Grant, 2001), and the movements of the mouth are synchronized with, or slightly precede changes in the auditory speech (Chandrasekaran et al., 2009; Vatakis et al., 2012). Temporal coherence has been implicated as a binding cue using non-speech stimuli (Atilgan et al., 2018; Atilgan & Bizley, 2020; Maddox et al., 2015) and can reduce effects of a masking stimulus in a speech detection task (Grant, 2001; Grant & Seitz, 2000).

Next, articulatory correspondence derives from the fact that acoustic speech sounds, phones, are linked with the specific mouth shape used to create them. Mouth shape provides information that is complementary to acoustic information, helping listeners to disambiguate place and manner of articulation (Grant & Bernstein, 2019; Grant & Walden, 1996). In order to benefit from this visual information, the correspondence of articulatory shape with specific phones must be learned based on experience with speech. We therefore theorized that higher order computations are necessary in order to benefit from integration of articulatory information in audiovisual speech.

Finally, talker congruence relates to whether the face and the voice originate from the same talker. This is also a learned correspondence based on experience with different individuals, both in a generalized sense (e.g., the average pitch of male talkers is lower than that of female talkers) and in a specific sense (e.g., creating associations between a known individual’s voice and face.)

Of these three cues, we hypothesized that temporal coherence would have the largest effect on performance based on previous work. Although in Fiscella et al. (2022) we did not see an effect of articulatory correspondence, we hypothesized that it could have a small effect with these stimuli where we could control articulatory content of the stimuli more completely. Finally, we expected to see no impact of talker identity congruence on binding under the assumption that complex processing of the voice and face would be required and not accessible to early regions performing binding computations.

### Experimental Design

The challenge of creating an experiment that is based on naturalistic speech stimuli, manipulates potential binding cues independently, and allows for subjects to make determinations about orthogonal features is not trivial. Here we created stimuli based on recordings of native English speakers smoothly transitioning between two vowels. These recordings then allowed us to create an auditory or visual representation of any vowel along that trajectory, modulated by a given timecourse. By selecting from a set of recordings representing different vowel pairs and two talkers, we were also able to manipulate articulatory and talker identity correspondence independently.

In a trio of experiments, we evaluated the relative contributions of temporal coherence (timecourse), articulatory correspondence (vowel pair), and talker identity correspondence (talker). In each experiment, we engaged participants in a dual task in which they responded as quickly and as accurately as possible to brief pitch deviations in the target auditory stream, as well as pale blue flashes over the lips of the video. This design emulates as many aspects as possible from Maddox et al. (2015). In the first experiment, we verified that binding was occurring for our artificial speech-like stimuli by considering two conditions: one in which the video matched the auditory target in all aspects (timecourse, vowel pair, and talker) and one in which the video instead matched the auditory masker. Subjects showed a clear benefit to performance when viewing the video that matched the auditory target. We then performed two further experiments to investigate the relative contributions of talker identity and articulatory congruence. In these experiments we saw no influence of the vowel pair matching, but we did identify a large impact of the talker in the video matching the auditory target. In fact, the combined results across all three experiments show that the identity of the talker in the video is the only factor that significantly influences performance in this task with these stimuli. Our findings, when considered alongside previous results, paint a more detailed picture of temporal coherence as a binding cue that specifically facilitates segregation of similar stimuli, while other cues such as talker congruence may aid selection.

## Experiment 1 Methods

All methods were preregistered: DOI 10.17605/OSF.IO/V8HTE.

### Subjects

We recruited 30 subjects (22 female, 8 male) between the ages of 18 and 40 (mean 23.7 ± 3.5 years). All subjects had normal hearing (thresholds of 20 dB or better for octave frequencies 500 Hz to 8 kHz) and self-reported normal or corrected-to-normal vision. Subjects gave informed consent and were paid for their participation. Of the 30 subjects, only 24 were included in our analysis. Five subjects did not pass the training described below and one did not meet our pre-defined d’ threshold of 0.8 during the main part of the experiment.

A power analysis using pilot data from the two authors (which could not be used in the final experiment as that are overtrained in the task and not naive participants) indicated that we would achieve sufficient power even with only 5 subjects. However, we expected that naive subjects will have more variable performance in their overall ability to do the task and in their alertness throughout data collection. As a more conservative decision, we based our sample size off of the original Maddox et al. (2015) paper showing the effect of temporal coherence. In that paper each task employed 16 subjects. We took that number and increased it by 50%.

### Equipment

Sounds were presented at 50 dB SPL and 48 kHz sampling rate by an RME soundcard through a KEF E301 free-field speaker at 0° azimuth and elevation. Visual stimuli were presented using an HTC Vive Pro Eye with a refresh rate of 90 frames per second. Responses were collected using the Vive controllers.

### Task

In each trial, subjects were presented with two auditory streams and one visual stimulus. The target auditory stream was present at the beginning of the trial and the masker stream was silent for the first three seconds. Subjects were asked to attend only to the first auditory stream (target) and visual stream and respond with a button press when they detected short auditory pitch changes in the voice of the target talker and visual blue flashes over the lips of the talker. They were instructed to ignore pitch events in the masker talker.

### Stimuli

We built the stimuli from audio-video recordings of two native English talkers (authors MSC and RKM) smoothly oscillating between a pair of two English vowels (/α/, /ε/, /i/, /o/, /u/) and additional audio recordings of the two talkers speaking a neutral vowel (sometimes called “schwa”) with even pitch and a duration of at least 14 seconds. A section of the videos that contained a clean vowel transition, no blinks, and minimal head movement was selected for further processing. Praat was used to extract the formants (F1, F2, and F3) of the vowel transitions. A monotonic regression was performed on each formant contour. Then time was warped such that the F1 transition was linear on the bark frequency scale with the endpoints being designated as 0 and 1. Interpolated formants of the linear model could then be applied to the neutral vowels using source-filter processing to synthesize any intermediate vowel sound between the original vowels in the pair. Video frames from the appropriate time stamp could be paired with this synthesized speech. Five vowel pairs were selected from a possible ten combinations as those with clear visual transitions for both talkers (1) /α/ + /o/, 2) /α/ + /u/, 3) /i/ + /u/, 4) /ε/ + /u/, 5) /ε/ + /o/).

For each frame, the eye position in the frame was detected by the dlib python package. These positions were smoothed using an averaging filter with a Hann window of width 41. With the smoothed eye positions, we used the imutils python package (2015/2022) to align and crop the image such that it was a 1024 x 1024 pixel square with the face centered, eyes level, and size normalized between the two talkers. The scale factor for each set of images was set using the first frame and applied to the rest of the set; however, each rotation and translation was based on the smoothed eye position for the given frame.

Each trial was 14 s in duration. The vowel identity was modulated by randomly generated timecourses. The timecourses were noise generated in the frequency domain with uniform energy between 0 and 7 Hz. Because talker RKM has choral experience, his overall vowel trajectories are more extreme than talker MSC. To compensate for this, we limited the timecourses applied to his vowels from 0.1 to 0.9 (rather than 0 to 1 for talker MSC). We additionally compensated for the discrepancies in the two talkers by accentuating talker MSC’s vowels. We applied the following nonlinearity twice to MSC’s timecourses:

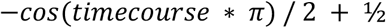

Pitch events were added to the existing pitch contour of each auditory stream using Praat. Each of these events was a 200 ms sinusoidal modulation with a 2-semitone depth. Visual events were a color change of the lips of the talker that lasted 200 ms. The color change was generated by overlaying a transparent blue mask over each frame during the 200 ms event. For each frame we first detected the lips using facial landmarks with the dlib python package. A transparent background was generated with the area of the lips shaded in opaque cyan (RGB: [0, 255, 255]). A Gaussian blur with kernel size of 33 x 33 pixels was applied to smooth the edges. The overlay was then added to the original video frame with an alpha of 0.2.

Event start times could only occur after the first 4 s of the trial (1 s after the onset of the masker stimulus), and before the last 1.5 s. Events in any stream were at least 1.2 s apart. In a given trial there were between three and five events distributed such that each stream had at least one event and up to three events.

### Experiment Structure

Prior to the main experiment, subjects completed a training module to slowly introduce the task (three short auditory-only trials with a single pitch event, three short visual-only trials with a single visual event, two full-length auditory only trials, four full-length audiovisual trials with no masker, and finally ten full trials). If subjects did not achieve an overall auditory sensitivity of 0.7 or higher and a visual hit rate of 0.8 in the training portion of the experiment, they were offered one opportunity to redo the training. Subjects who could not meet these criteria on the second try were excused from participation.

There were two conditions in the main experiment, one in which the video matched the audio of the target talker and one in which the video matched the audio of the masker talker. There were a total of 80 trials in the experiment across the video match conditions and counterbalancing of stimuli (2 conditions x 2 target talkers x 5 target vowel pairs x 4 masker vowel pairs). Participants were offered a self-timed break every ten trials. Each session was approximately 45 minutes to 1 hour in duration. Any participant who did satisfy the same criteria of an overall auditory sensitivity of 0.7 or higher and a visual hit rate of 0.8 in the main experiment was excluded from analysis.

### Analysis

We measured the sensitivity (d’) of the auditory task for each condition. A hit was defined as a response within 1 s of a pitch event in the target talker. A false alarm was a response within 1 s of a pitch event in the masker talker. A miss was no response with within 1 s of a pitch event in the target talker. A correct rejection was no response within 1 s of a pitch event in the masker talker. We also calculated the hit rate for visual events across the whole experiment. Visual hit rate was defined as the number of visual events for which there is a response within 1 s over the total number of visual events.

### Statistical Model

To gain a holistic view of each variable’s contribution to task performance, we fit a generalized linear model to the response data. Each auditory event (target, coded as a 1, or masker, coded as a 0) was one data point in the model. Event type (whether the event was in the target or masker stream, coded as 1 and 0 respectively), video match (of the event stream with the video, coded as 1 for matching and 0 for not matching), and the interaction of event type and video match were fixed effects in the model. We also included random intercepts for the talker of the event stream, the vowel pair of the event stream, their interaction, and the vowel pair of the other stream. Finally, we included a random intercept for each subject as well as random slopes for event type and video match, the maximal structure such that the model still converged.

## Experiment 1 Results

We analyzed data from all 24 subjects who clearly exceeded our data-inclusion thresholds (Figure 2). The main indicator of auditory performance, sensitivity (d’), improved when the auditory target stream matched the video (Figure 2A). This performance improvement was accompanied by a slight increase in bias (Figure 2B). By considering hits (Figure 2C) and false alarm rates (Figure 2D) separately, the improvement can be seen to be driven mainly by a decrease in false alarm rate. All subjects performed close to ceiling in the visual task, confirming they were attending the video.

**Figure 1:**
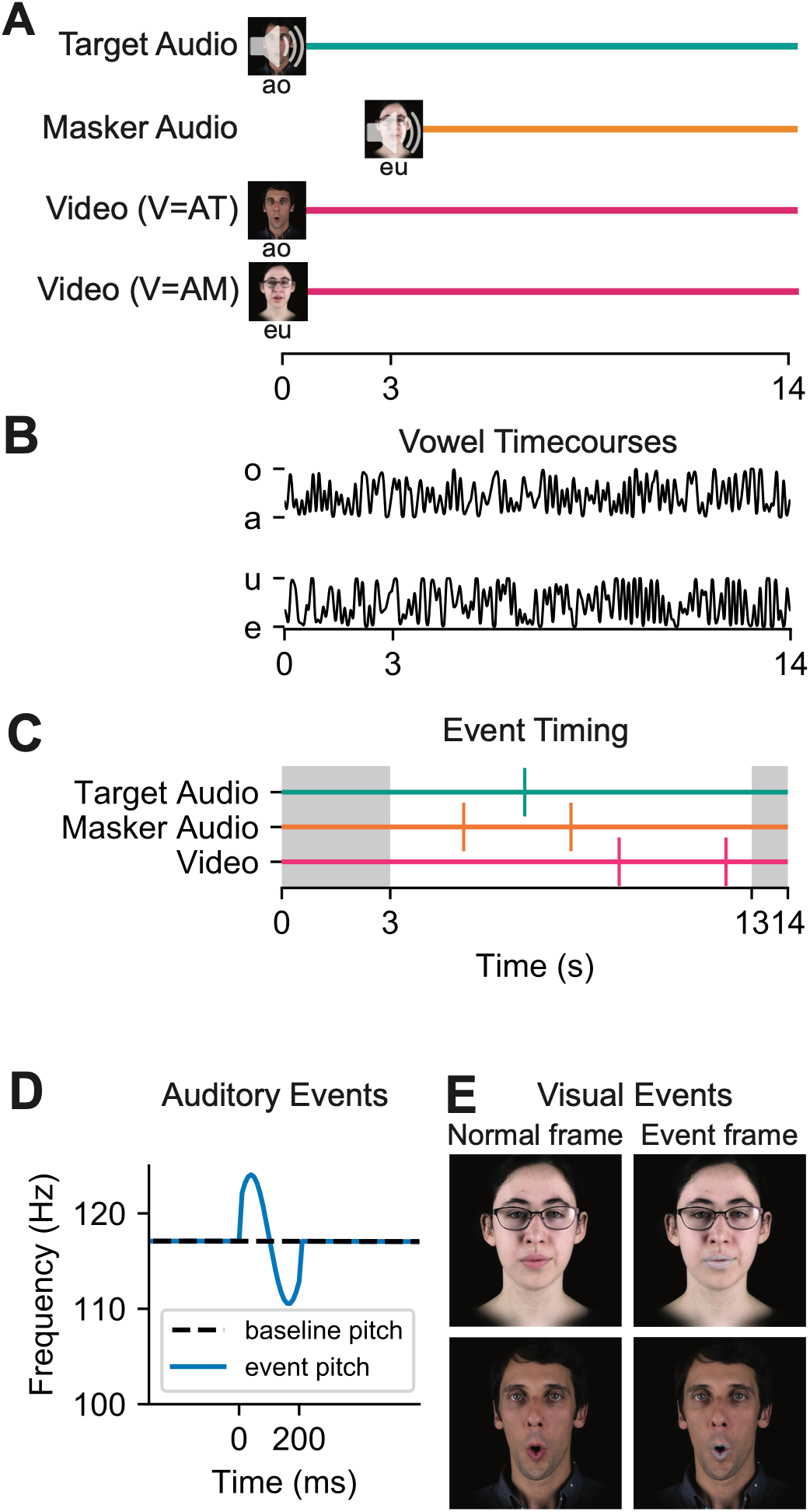
Task explanation. A) Timing of stimuli. Target and masker audio are shown with their two potential visual counterparts. B) Example timecourses. C) Example event times. Vertical lines show the timing of an event. D) The pitch trajectory of an auditory event. E) The two talkers in both normal frames and event frames. [Identifiable images depict authors MSC and RKM.]

**Figure 2:**
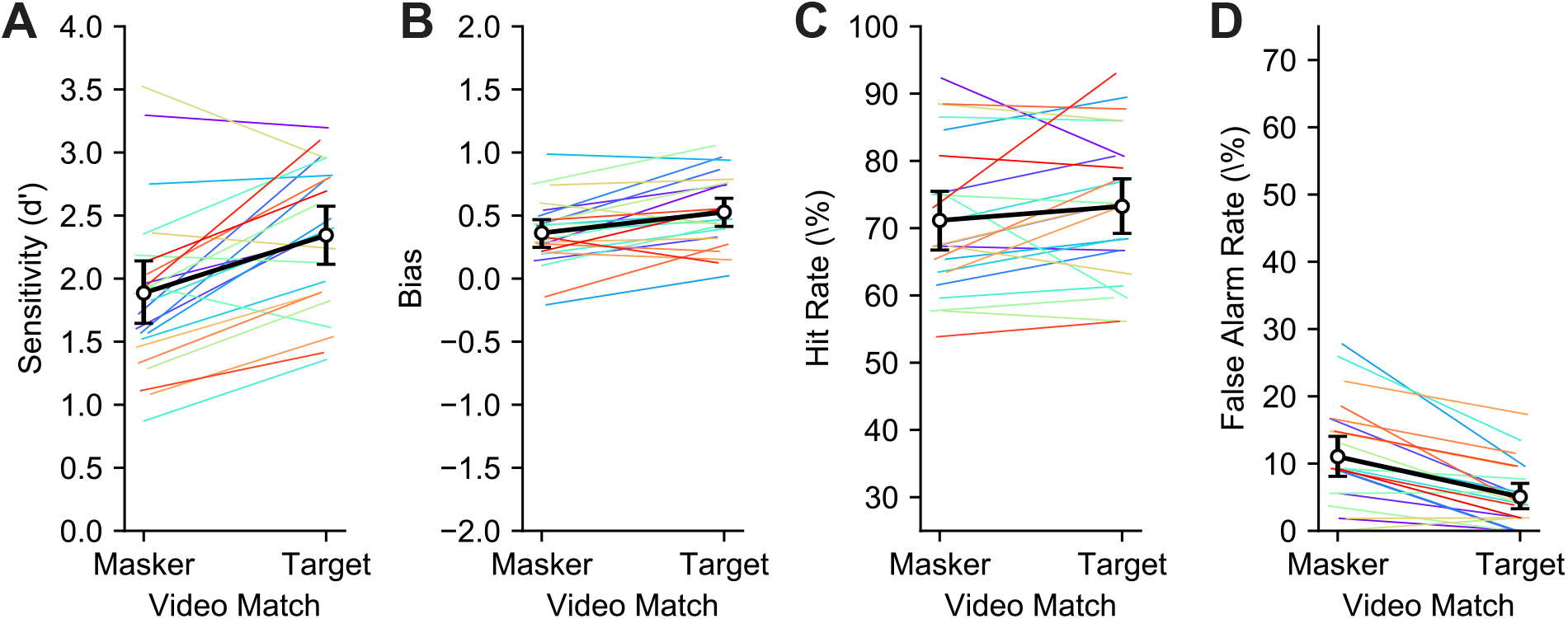
Auditory performance improves when the auditory target matches the video. Thin colored lines indicate single subjects. Thicker black lines indicate the mean across subjects. Error bars indicate 95% confidence intervals based on resampling subjects with replacement. A. Auditory sensitivity (d’). B. Bias, where a positive bias indicates a bias towards not responding (false alarms considered more costly than misses). C. Hit rates. D. False alarm rates.

Our qualitative observations are borne out in the statistical model (Table 1). We saw significant effects of whether the event was in the target or masker stream, whether the stream containing the event matched the video, and their interaction. The large positive model term for event type (i.e., target or masker) indicated that participants’ behavior was driven by the task (2.53, **p<2×10^-16^**). Subjects should respond to events in the target stream (hits) but not in the masker stream (false alarms). The effect of video match showed that a coherent visual stream significantly improved performance (0.500, p=6.35e-10) either by increasing hits or decreasing false alarms. Finally, the interaction term indicated that the benefit of a matching visual stream was driven by a reduction in false alarms (−0.527, **p=1.57×10^-7^**).

To explain this interpretation, Figure 3 presents a visual representation of the model terms. There are four types of events in the experiment: two video match trial conditions and two auditory streams that can contain an event. The video match term in the statistical model does *not* correspond to the trial type. Instead, video match is 1 when there is a target event in a video match target (V=AT) trial *and* when there is a masker event in a video match masker (V=AM) trial. The effect of the video match term in the statistical model is to assign whether responses increase or decrease for a matched video. In the first column of Figure 3, which shows a positive video match term, responses increase in the blue boxes (video match) and decrease in the grey boxes (video mismatch). In the second column, the opposite is true. Each row of the figure shows a different sign of the interaction term. In the first row, showing a positive interaction of video match and event type, the interaction term indicates a stronger effect of video match when the event type is target. This means that responses to target events (hit rate) drive the change in performance across trial types. In the third row, showing a negative interaction, the opposite is true. There is a stronger effect of video match when the event type is masker (i.e., *not* target), so we see the change in performance dominated by false alarms. Although we only show a visual explanation of the statistical model for Experiment 1, the same logic applies to Experiments 2 and 3.

**Figure 3:**
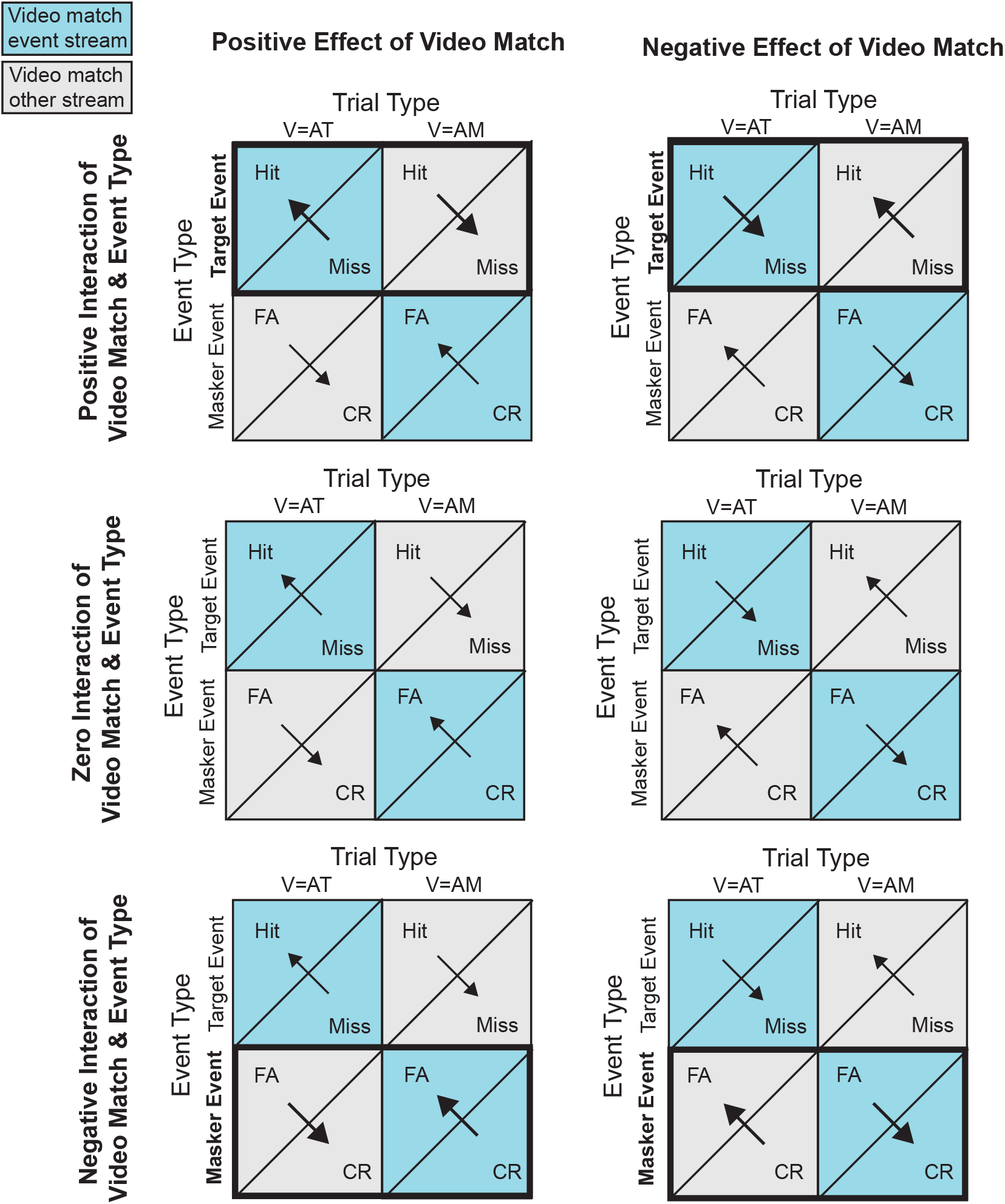
Explanation of statistical model terms. Columns and rows indicate possible effect directions. Each grid shows the possible outcomes for the four types of events in the experiment (2 trial type x 2 event type). Arrows show the direction of change based on the effect of video match with their weight and size indicating larger or smaller changes. Thick boxes indicate interactions, specifically that there is a stronger effect of video match for one event type.

## Experiment 2 Methods

We followed the methods from Experiment 1 with the following deviations. All methods were preregistered: DOI 10.17605/OSF.IO/BCFWZ

### Subjects

Thirty-three subjects (25 female, 8 male, age 24.4 ± 3.3 years) participated in Experiment 2. Twenty-three of these subjects had also participated in Experiment 1.

In our first experiment we recorded 40 trials/condition for 24 subjects. Our data showed that this sample size is in excess of that required to show an effect of overall congruence across all subjects. Here we expected the effect of talker congruence to be smaller, but had no preconception of how much smaller. Thus, we chose a sample size that was almost certain to replicate the preliminary effect from experiment 1 and at the upper limit of what is feasible given numerous practical constraints on human subject data collection. In experiment 1, 40 trials * 24 subjects gave us 960 trials per condition. In this experiment, we had 30 trials/condition, which required we record from 32 subjects to get the same 960 trials/condition across subjects.

### Experimental Conditions

Previously, auditory streams could match the video in all or none of the following aspects: timecourse, vowel pair, and talker. In Experiment 2, we separated timecourse and vowel pair from talker. In each trial, the target or masker auditory timecourse and vowel pair could match the video timecourse and vowel pair (V_timecourse/vowel_ = AT_timecourse/vowel_ or V_timecourse/vowel_ = AM_timecourse/vowel_ respectively), and the target or masker auditory talker could match the video talker (V_talker_ = AT_talker_ or V_talker_ = AM_talker_ respectively). The condition in which V_timecourse/vowel_ = AT_timecourse/vowel_ or V_timecourse/vowel_ and V_talker_ = AT_talker_ is the same as V=AT from Experiment 1 and V_timecourse/vowel_ = AM_timecourse/vowel_ and V_talker_ = AM_talker_ is the same as V=AM.

The main part of Experiment 2 included 160 trials (2 timecourse/vowel match x 2 talker match x 2 target talkers x 5 target vowels x 4 masker vowels).

### Statistical Model

We fit a generalized linear model as before with the following fixed effects: event type (whether the event was in the target or masker stream, coded as 1 and 0 respectively), talker match (of the event stream with the video, coded as 1 for matching and 0 for not matching), timecourse/vowel-pair match (following the same rules as talker match), the interaction of event type with talker match, the interaction of event type with timecourse/vowel-pair match, and the three-way interaction of event type, talker match and timecourse/vowel-pair match. As before, we included random intercepts for the talker of the event stream, the vowel pair of the event stream, their interaction, and the vowel pair of the other stream. We included a random intercept for each subject as well as a random slope for the event type. No other random effects could be included without impacting model convergence.

## Experiment 2 Results

All subjects who passed the training module performed the auditory and visual tasks well above chance. Subjects increased their performance when the auditory target talker matched the video talker (V_talker_ = AT_talker_ vs. V_talker_ = AM_talker_), especially when the auditory timecourse and vowels matched the video (V_talker_ = AT_talker_, V_timecourse/vowel_ = AT_timecourse/vowel_) (Figure 4A). There was also a modest decrease in performance when the auditory timecourse and vowels matched the video (V_timecourse_ = AT_timecourse_ vs. V_timecourse_ = AM_timecourse_). Subjects were generally biased towards not responding (more misses than false alarms), and there appears to have been a decrease in bias in the V_talker_ = AM_talker_, V_timecourse_ = AT_timecourse_ condition (Figure 4B). Hit rate (Figure 4C) was relatively stable compared to false alarm rate (Figure 4D).

**Figure 4:**
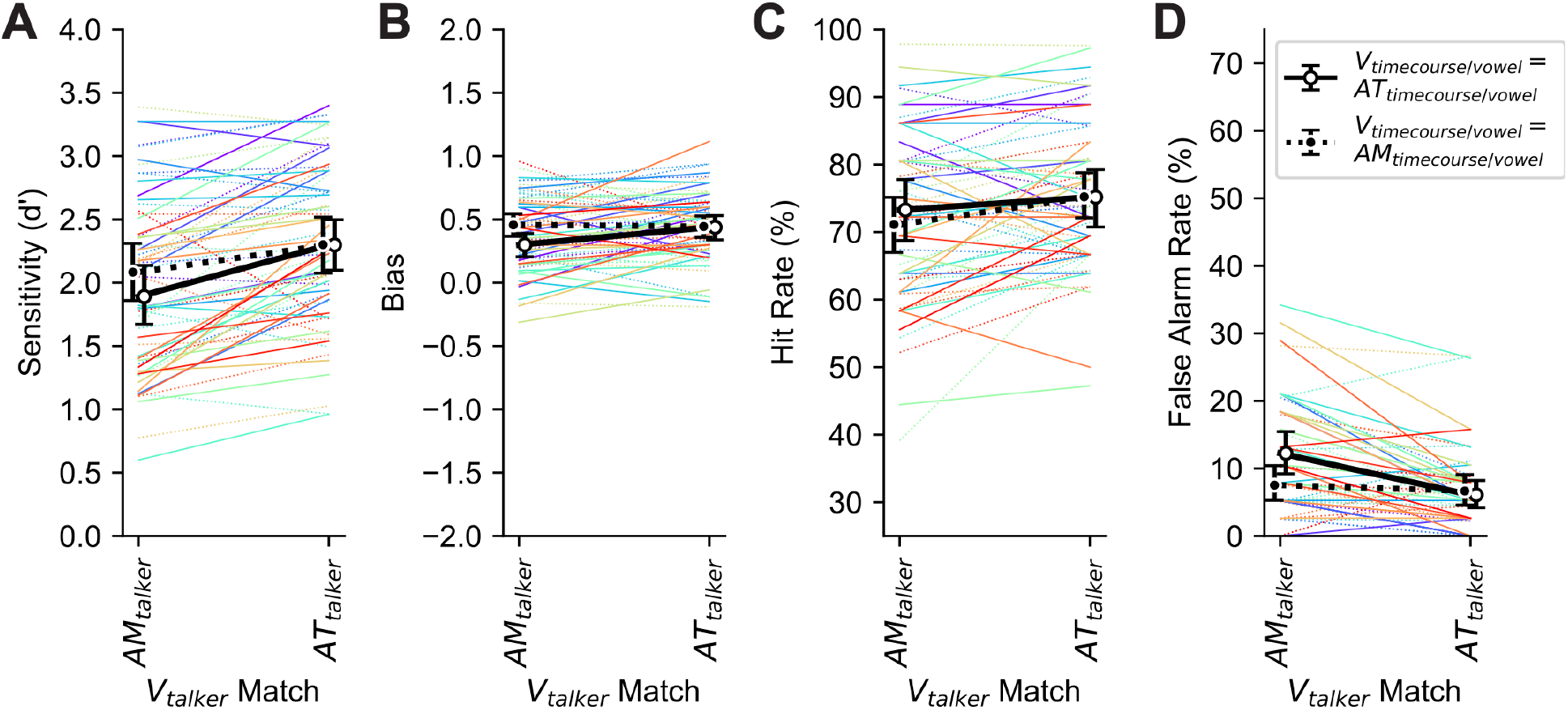
Auditory performance improves when the auditory target talker matches the video, especially if the target stream’s timecourse and vowel pair matches the video. Thin colored lines indicate single subjects. Thicker black lines indicate the mean across subjects. Error bars indicate 95% confidence intervals based on resampling subjects with replacement. A. Auditory sensitivity (d’). B. Bias. C. Hit rates. D. False alarm rates.

We fit the data to the generalized linear mixed effects model that we described in our preregistration (Table 2). A separate model, including the interaction term between temporal coherence and talker congruence that is needed to better describe the data, is presented in a later section describing unregistered analyses. In this model, we again saw clear evidence of subjects performing the task as instructed from the event type term (2.25, **p<2×10^-16^**). The vowel and timecourse match term showed a modest decrement to performance (−0.17, **p=0.00188**), while talker match benefitted performance (0.269, **p=7.50×10^-7^)**. There was a positive, significant interaction between whether the event is in the target stream and whether the event vowel and timecourse match the video (0.245, **p=0.00157**). This positive interaction term reflects the decrease in bias, and thus a larger decrease in misses relative to the increase in false alarms for V_timecourse/vowel_ = AT_timecourse/vowel_ trials. The interaction of event type and talker match in this model was not significant. In the unregistered analysis section, the model which includes the missing interaction term showed a significant negative interaction of event type and talker match.

**Table 1:**
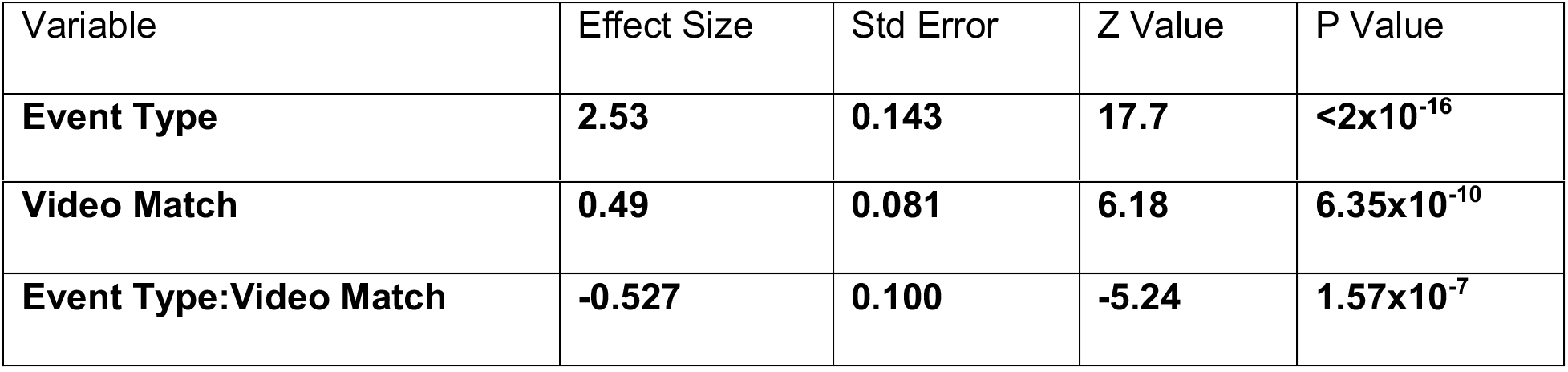
GLM fit of Experiment 1 data.

| Variable | Effect Size | Std Error | Z Value | P Value |
| --- | --- | --- | --- | --- |
| **Event Type** | **2.53** | **0.143** | **17.7** | **$<2 \times 10^{-16}$** |
| **Video Match** | **0.49** | **0.081** | **6.18** | **$6.35 \times 10^{-10}$** |
| **Event Type:Video Match** | **-0.527** | **0.100** | **-5.24** | **$1.57 \times 10^{-7}$** |

**Table 2:** GLM fit of Experiment 2 data as specified in preregistration.

| Variable | Effect Size | Std Error | Z Value | P Value |
| --- | --- | --- | --- | --- |
| **Event Type** | **2.25** | **0.131** | **17.4** | **$< 2 \times 10^{-16}$** |
| **Timecourse/Vowel Match** | **-0.169** | **0.0544** | **-3.109** | **0.00188** |
| **Talker Match** | **0.269** | **0.043** | **4.95** | **$7.50 \times 10^{-7}$** |
| **Event Type:**<br>**Timecourse/Vowel Match** | **0.245** | **0.0778** | **3.16** | **0.00157** |
| Event Type:Talker Match | -0.123 | 0.0763 | -1.61 | 0.108 |
| Event Type:Talker Match:<br>Timecourse/Vowel Match | -0.0511 | 0.0805 | -0.634 | 0.526 |

## Experiment 3 Methods

We followed the methods from Experiment 1 with the following deviations.

All methods were preregistered: DOI 10.17605/OSF.IO/AKRVD

### Subjects

Thirty-three subjects (25 female, 8 male, age 24.0 ± 3.9 years) participated in Experiment 3. Twenty subjects participated in both Experiment 1 and 2. Eight had participated in Experiment 2 but not Experiment 1. One subject was excluded from analysis because their overall performance fell below our threshold.

We again aimed for 32 subjects for the same justification listed in Experiment 2.

### Experimental Conditions

In Experiment 3, we investigated the effect of vowel pair independent of timecourse and talker. In each trial, the target or masker auditory timecourse and talker could match the video timecourse and talker (V_timecourse/talker_ = AT_timecourse/talker_ or V_timecourse/talker_ = AM_timecourse/talker_ respectively). Additionally, the target or masker auditory vowel pair could match the video fully or not at all *or*, in the case where the target and masker vowel pairs shared one vowel, there could be a partial match. For the case in which the target and masker acoustic vowel pairs shared no vowels, there were two conditions: the video had a vowel identity that is fully congruent with the target auditory stream (V_vowelpair_ =AT_vowelpair_), or was fully congruent with the masker auditory stream (V_vowelpair_ =AM_vowelpair_). When the target and masker acoustic vowel pairs shared one vowel, we had two additional conditions: fully congruent with the target auditory stream and sharing one vowel with the masker auditory stream (V_vowelpair_ =AT_vowelpair_ ~AM_vowel_), or fully congruent with the masker auditory stream and sharing one vowel with the target auditory stream (V _vowelpair_ =AM_vowelpair_ ~AT_vowel_).

The main part of Experiment 3 included 160 trials selected from a possible 320 trials (4 vowel match x 2 timecourse/talker match x 2 target talkers x 5 target vowels x 4 masker vowels). Each subject was presented with a different random selection of trials.

### Statistical Model

We fit a generalized linear model as before with the following fixed effects: event type (whether the event was in the target or masker stream, coded as 1 and 0 respectively), vowel match (of the event stream with the video, coded as 2 for two matching vowels, 1 for one matching vowel and 0 for no matching vowels), timecourse/talker match, the interaction of event type with vowel match, the interaction of event type with timecourse/talker match, and the threeway interaction of event type, vowel match and timecourse/talker match. As before, we included random intercepts for the talker of the event stream, the vowel pair of the event stream, their interaction, and the vowel pair of the other stream. Finally, we included a random intercept for each subject as well as a random slope for the event type.

## Experiment 3 Results

Subjects showed a modest improvement in d’ when the timecourse and talker of the target auditory stream matched the video (Figure 5A). This improvement was paired with an increase in bias (Figure 5B), a relatively stable hit rate stable (Figure 5C), and a lower false alarm rate in V_timecourse/talker_=AT_timecourse/talker_ conditions (Figure 5D). There was little effect of vowel match on any measure of task performance. Performance in the visual task was consistently high.

**Figure 5:**
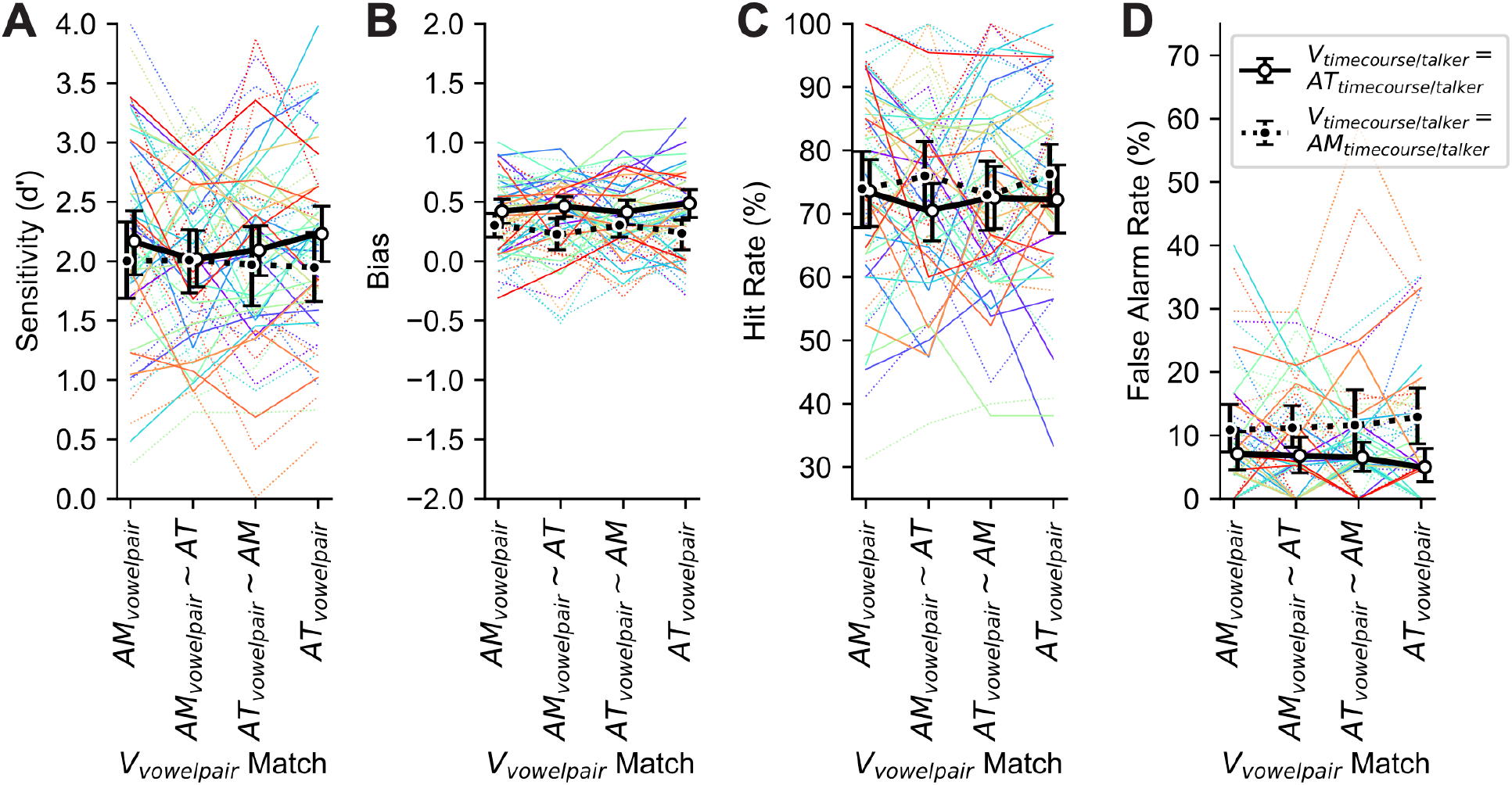
Performance improves when the video talker and timecourse match the target auditory stream. Thin colored lines indicate single subjects. Thicker black lines indicate the mean across subjects. Error bars indicate 95% confidence intervals based on resampling subjects with replacement. A. Sensitivity. B. Bias. C. Hit rate. D. False alarm rate.

**Figure 6:**
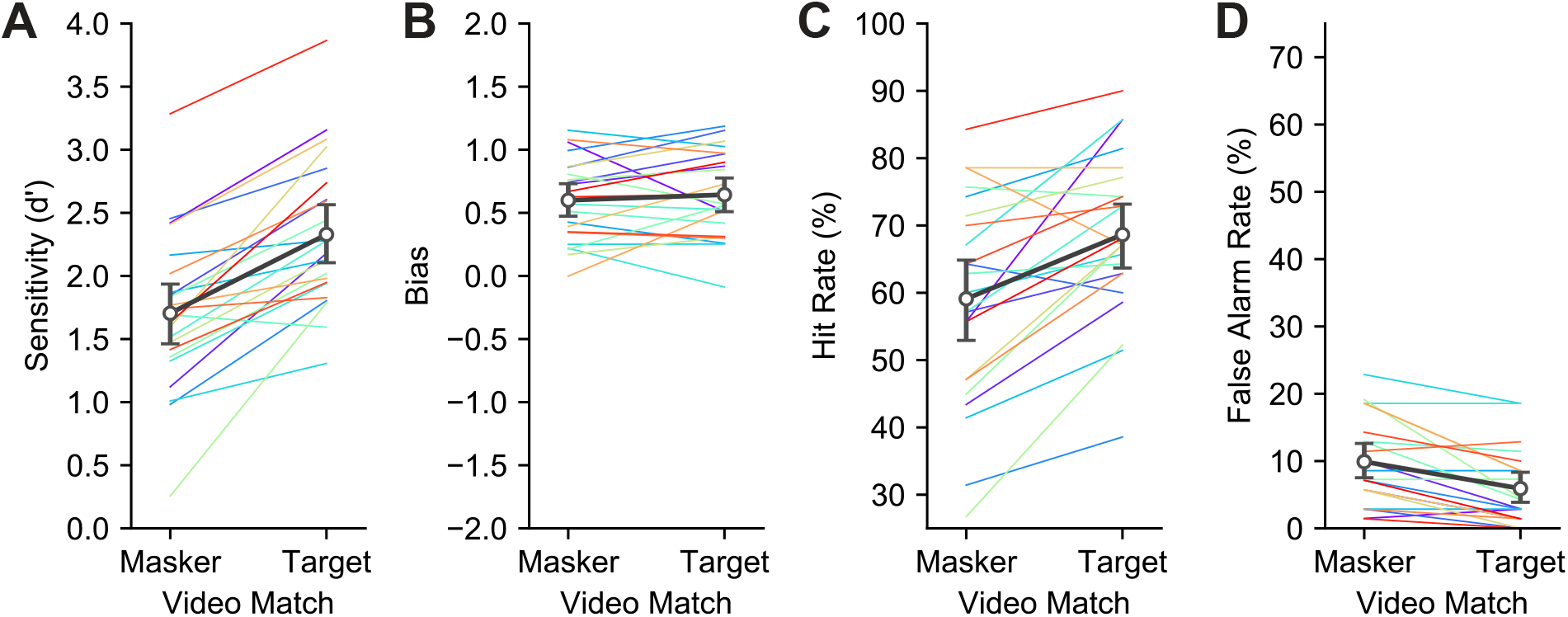
Data from Fiscella et al. (2022) reanalyzed to examine the specific effects of talker match on task performance. Here we present averages across orientation conditions and solely examine the difference between the target and masker video being presented. Dark lines/large markers show across subject averages while thin colored line indicate individual subject data. Error bars show 95% confidence intervals based on resampling subjects with replacement. A. Sensitivity. B. Bias. C. Hit rate. D. False alarm rate.

Again, we fit the data to a generalized linear mixed effects model described in our preregistration. The version of the model with the interaction term between timecourse/talker and vowel match is presented in the following section. In the preregistered model we saw performance in the task is good (2.57, p<**2×10^-16^**). Timecourse and talker match was associated with a clear increase to performance (0.35, p=**2.81×10^-10^**). Vowel match offered no significant benefit (0.02, p=0.51). There was a negative, significant interaction between whether the event was in the target stream and whether the event timecourse and talker matched the video (−0.34, p=**0.00057**). This is the same case discussed in Experiment 1 where the benefit created by viewing a video that matched the timecourse and talker of the target audio benefitted performance by reducing false alarms. The remaining interaction terms were not significant.

## Unregistered Analysis Methods

Both Experiments 2 and 3 were preregistered with a model structure that excluded the interaction term between the two match conditions. For each, we fit an additional generalized linear mixed effects model that was identical to the preregistered model except for an added interaction of timecourse/vowel match with talker for Experiment 2 and interaction of timecourse/talker with vowel match for Experiment 3.

We also fit a model to the data from all three experiments to determine which effects were strongest across experiments. The included fixed effects for that overall model were event type, timecourse match, talker match, and vowel match. We also included interaction terms between event type and each of the three match variables. Finally, we included random intercepts for the talker and vowel pair of the event stream, as well as their interaction. Subject identity was also considered a random intercept and we fit a random slope of session with subject to account for any learning or difference between experiment difficulty.

Finally, we performed additional analysis of data previously published in Fiscella et al (2022) for comparison with the present results. The experiment included two video match conditions (video match target and video match masker, as in Experiment 1) and two video orientation conditions (upright and inverted). We calculated sensitivity, bias, hit rate, and false alarm rate for video match conditions, averaging across orientations. We also fit the data to a linear model that was as similar as possible to those fit to Experiments 1–3. The model was fit on the level of each trial (as this was a discrimination task rather than a detection task) where response was predicted by the following fixed effects: trial type, video match, video orientation, and interactions of video match with video orientation, trial type with video match, trial type with video orientation, and the three-way interaction of trial type, video match, and video orientation. A random intercept for subject was also included.

## Unregistered Analysis Results

Table 4 shows a model of Experiment 2 that includes the interaction of timecourse/vowel match with talker match. Consistent with observations from Figure 4, there is no consistent role of timecourse/vowel match on its own, but there is a significant interaction with talker match (−0.37, p=**0.00055**). This negative interaction indicates that a matching timecourse and vowel pair only hinders performance when the target auditory talker does not match the video or, equivalently, that the effect of talker match is stronger when the auditory target timecourse and vowel pair match the video. There is a positive three-way interaction of event type, and both match variables (0.33, p=**0.016**) which represents both a more dramatic change in false alarms when the video’s timecourse/vowel match the auditory target and a slight increase in hit rate across talker match conditions when the video timecourse/vowel match the masker auditory stream. We see no overall effect of the timecourse and vowel pair matching the video in this model (0.03, p=0.65).

**Table 3:** GLM fit of Experiment 3 data as specified in preregistration.

| Variable | Effect Size | Std Error | Z Value | P Value |
| --- | --- | --- | --- | --- |
| **Event Type** | **2.57** | **0.267** | **9.61** | **$<2 \times 10^{-16}$** |
| **Timecourse/Talker Match** | **0.355** | **0.0563** | **6.31** | **$2.81 \times 10^{-10}$** |
| Vowel Match | 0.0217 | 0.0329 | 0.660 | 0.509 |
| **Event Type:**<br>**Timecourse/Talker Match** | **-0.336** | **0.0976** | **-3.45** | **0.000566** |
| Event Type:Vowel Match | -0.0115 | 0.0497 | -0.232 | 0.817 |
| Event Type:Vowel Match:<br>Timecourse/Talker Match | -0.0383 | 0.0527 | -0.726 | 0.468 |

**Table 4:** Experiment 2 GLM including the interaction of timecourse/vowel match with talker match.

| Variable | Effect Size | Std Error | Z Value | P Value |
| --- | --- | --- | --- | --- |
| **Trial Type** | **2.36** | **0.135** | **17.4** | **$<2 \times 10^{-16}$** |
| Timecourse/Vowel Match | 0.036 | 0.081 | 0.459 | 0.65 |
| **Talker Match** | **0.453** | **0.077** | **5.90** | **$3.66 \times 10^{-9}$** |
| **Timecourse/Vowel Match:Talker Match** | **-0.378** | **0.109** | **-3.45** | **$5.5 \times 10^{-4}$** |
| Trial Type:<br>Timecourse/Vowel Match | 0.0407 | 0.0979 | 0.415 | 0.67 |
| **Trial Type:Talker Match** | **-0.306** | **0.0936** | **-3.274** | **0.0011** |
| Trial Type:Talker Match:<br>Timecourse/Vowel Match | 0.327 | 0.136 | 2.41 | 0.0162 |

By including the interaction of timecourse/talker and match vowel match (Table 5), we resolved a marginal, but insignificant effect of vowel match (0.1, p=0.051). The interaction term, which was also marginally significant then clarifies that any effect of vowel match is present primarily when the target timecourse and talker match the video (−0.13, p=0.044). Otherwise, this term did not change the significance of any model parameters.

**Table 5:** Experiment 3 GLM including the interaction of timecourse/talker match with vowel match.

| Variable | Effect Size | Std Error | Z Value | P Value |
| --- | --- | --- | --- | --- |
| **Event Type** | **2.67** | **0.273** | **9.81** | **$<2 \times 10^{-16}$** |
| **Timecourse/Talker Match** | **0.532** | **0.105** | **5.07** | **$4.03 \times 10^{-7}$** |
| Vowel Match | 0.101 | 0.0516 | 1.951 | 0.051 |
| **Timecourse/Talker Match:Vowel Match** | **-0.135** | **0.0669** | **-2.01** | **0.044** |
| **Event Type:**<br>**Timecourse/Talker Match** | **-0.514** | **0.132** | **-3.90** | **$9.78 \times 10^{-5}$** |
| Event Type:Vowel Match | -0.0905 | 0.0636 | -1.42 | 0.155 |
| Event Type:Vowel Match:<br>Timecourse/Talker Match | 0.0965 | 0.0852 | 1.13 | 0.257 |

When analyzing data from all three experiments in a single model (Table 6) there was a clear significant effect of the talker match (0.36, p<**2×10^-16^**) and a negative interaction of event type with talker match (−0.33, p=3.42e-10). Timecourse match and vowel match had no significant effect on performance across experiments, nor do their interactions with event type.

**Table 6:**
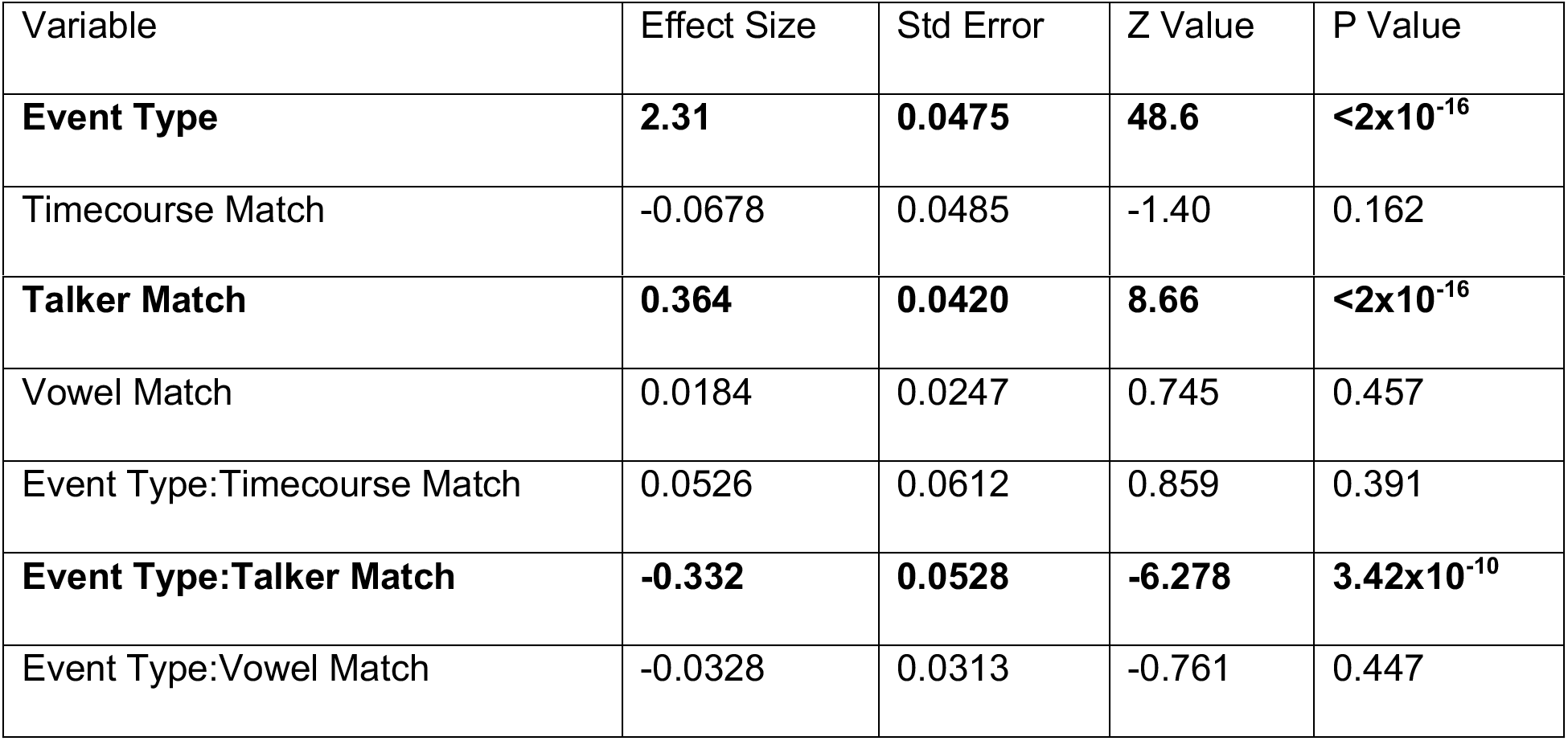
Model including all data from Experiments 1, 2, and 3.

### Analysis of previously collected data from Fiscella et al., 2022

We previously recorded data from listeners performing a similar task, but with full sentences and orthogonal modulation applied to the whole trial. Although we reported sensitivity in that paper, we did not report hit rate or false alarm rate. Here we reanalyze our previous data to look at what drove the change in sensitivity. Full methods are presented in the original paper.

Table 7 shows the GLM model of our previous data. In this experiment, only event type (1.81, p**<2×10^-16^**) and video match (0.244, p=**0.00813**) significantly impacted performance. Importantly, the interaction of event type and video match was near zero (−0.00894, p=0.934) indicating that neither hit rate nor false alarm rate dominated changes in performance.

**Table 7:** GLM fit of data from Fiscella et al (2022).

| Variable | Effect Size | Std Error | Z Value | P Value |
| --- | --- | --- | --- | --- |
| **Trial Type** | **1.81** | **0.0831** | **21.7** | **<2x10<sup>-16</sup>** |
| **Video Match** | **0.244** | **0.0921** | **2.65** | **0.00813** |
| Video Orientation | -0.154 | 0.103 | -1.50 | 0.134 |
| Video Match:Video Orientation | 0.107 | 0.135 | 0.796 | 0.425 |
| Trial Type:Video Match | -0.00894 | 0.113 | -0.079 | 0.937 |
| Trial Type:Video Orientation | 0.114 | 0.121 | 0.937 | 0.349 |
| Trial Type:Video Match:Video Orientation | -0.0515 | 0.163 | -0.326 | 0.752 |

## Discussion

### Overall summary

We performed three experiments investigating three potential contributors to audiovisual binding with speech-like stimuli and found that only the identity of the talker showed a significant and consistent effect on detection of orthogonal pitch events. Despite former research showing that the temporal coherence of audiovisual stimuli is sufficient to bind two streams and facilitate detection of orthogonal features, temporal coherence provided no measurable benefit to performance in our task. Across experiments, however, we saw a clear benefit to performance when the visual talker identity matched the talker voice in the auditory stream.

The benefit of viewing a matching talker was broadly carried by a reduction in false alarm rates. This implies that listeners were improving their ability to filter out the masker talker. In contrast, previous reports of binding for non-speech stimuli have shown benefits to sensitivity that are dominated by changes in hit rate (Maddox *et al*., 2015; Atilgan & Bizley, 2020). The large role of hit rate suggests that in previous studies, temporal coherence has improved the perception of the target auditory stream itself (rather than allowing the listener to “tune out” the masker stream). We also reanalyzed data previously presented in Fiscella et al. (2022) and found that the change in sensitivity was equally supported by an increase in hit rate and a decrease in false alarms. Differences in bias effects (related to the ratio of hits and false alarms) across experiments suggest that two separate mechanisms may be associated with changes in overall sensitivity.

### Audiovisual binding and selective attention

Audiovisual binding is defined as the early combination of auditory and visual stimuli such that an observer can form and attend to a cross-modal perceptual object (Bizley et al., 2016). The present task, similar to its predecessors (Atilgan & Bizley, 2020; Fiscella et al., 2022; Maddox et al., 2015), required observers to detect orthogonal features in both a visual stream and a target auditory stream while ignoring features in a masker auditory stream. If the observer binds the visual and target auditory streams, they can attend a single object rather than splitting their attention across two objects. The attentional benefit is then, canonically, measured as an improvement in sensitivity (d’). However, it is critical to note that selective attention comprises two stages: the first is to segregate the scene into perceptual objects (audio-visual binding), and the second is allocating attention to objects of interest (selection).

Conspicuously, the experiments presented here showed no evidence that temporal coherence benefits detection of orthogonal features. We propose that this finding is due to a fundamental difference in the ease of segregating auditory streams across this and former experiments. Segregation, the forming of discrete auditory objects or streams from a mixture of sounds, relies on several cues, including features like pitch, temporal envelope, and timbre (Bregman et al., 1990; Cusack & Roberts, 2000; Dannenbring & Bregman, 1976; Grimault et al., 2002; Singh, 1987; Singh & Bregman, 1997; Vliegen et al., 1999; Vliegen & Oxenham, 1999). Previous experiments that showed an effect of temporal coherence in audiovisual binding involved stimuli that were more difficult to segregate. In those using non-speech stimuli (Atilgan & Bizley, 2020; Maddox et al., 2015), the target and masker streams had only a modest difference in fundamental frequency (440 Hz vs. 565 Hz) and different envelopes. In Fiscella et al. (2022), there were a total of three masking voices, rendering the overall scene more complex. There was always at least one talker of the same sex, and therefore similar pitch, as the target talker in the scene. Here, segregation was aided by potent cues. The two voices were separated by a much larger relative difference in pitch (~100 Hz vs. ~200 Hz) and there was always a difference in timbre carried by the vowel identities. The masker talker in the current experiments could thus be easily segregated, so using temporal coherence to bind the visual stream with the target auditory stream offered no additional benefit to performance.

Given that temporal coherence did not benefit the listener in this task, it is then surprising that talker identity congruence offered such a consistent benefit. One possible interpretation is that seeing the face of the relevant talker offers a salient cue for maintaining selective attention on the correct talker, even without audiovisual binding. In other words, seeing the correct face makes it easier to keep track of who you are supposed to be listening to. This high-level benefit, independent of audiovisual binding, might explain why we saw a change in false alarm rate, rather than hit rate, in this task.

### The continuum of speech-like stimuli

Across many important and informative studies, numerous stimuli have been called “speech.” However, few speech stimuli, including those employed in this work, could be construed as natural, conversational verbal communication signals. To fully understand differences between the results of this study and of Fiscella et al. (2022) and to motivate future studies, it is crucial to discuss the differences between our stimuli and true natural speech.

First, natural speech involves both vowels *and* consonants. Our stimuli here represent a very limited subset of speech sounds, a selection of common English vowels. In auditory speech, the combination of consonants with vowels is important to their identification (Strange et al., 1976). The success of lipreading in audiovisual speech, a proxy for the quality of visual information, also depends on context and coarticulation (Benguerel & Pichora-Fuller, 1982; Vroomen & de Gelder, 2001). Therefore, information is lost by presenting vowels in isolation from consonants. Furthermore, models of vowel discrimination show that audiovisual information only offers a modest improvement, if any, to audio-only vowel information at 0 dB SNR (Teissier et al., 1997) where similar models describing consonant discrimination show larger benefits of audiovisual information at the same SNR (Grant & Walden, 1996). We also must consider the temporal dynamics of consonants and vowels. In general, speech sounds are preceded by visual articulatory features with an overall stimulus onset asynchrony (SOA) of 100–300 ms (Chandrasekaran et al., 2009). Many consonants require a preparatory mouth movement that allows the talker to produce the correct articulation. While vowels in natural speech may be preempted by a movement of the tongue prior to voicing (Alfonso & Baer, 1982), this comprises a shorter delay than for consonants and is only the case when there is a voicing onset. This difference in the temporal dynamics of speech production is also reflected in the temporal windows of integration for consonants and vowels (Vatakis et al., 2012). Here we used continuous vowels with no interruptions. These auditory vowels, constrained by the physics of sound production, are synchronous with the visual face used to articulate them. Generally, by limiting the representation of speech elements we compromise on our representation of speech, and risk failing to observe any effect of multisensory integration that relies on the dynamics of consonants or on the combination of consonants with vowels.

Second, natural speech has a temporal envelope characterized by the syllabic and prosodic structures of words and phrases. While our stimuli were modulated in terms of their timbre, their overall amplitude envelope was not modulated. Furthermore, the frequencies included in the timecourses (0–7 Hz) slightly exceeded normal syllabic rates, although syllabic rate is highly dependent on who is speaking and what they are saying (Coupé et al., 2019; de Johnson et al., 1979; Jacewicz et al., 2009). Diphthongs can be quite short (175 –300 ms), depending on the person speaking (Jacewicz et al., 2003), although this still leads to a maximum rate of 5.7 Hz, slightly below 7 Hz. If temporal coherence is an audiovisual binding cue for speech, it is possible that we did not measure an effect of timecourse here due to processing that is selective for amplitude modulation or a narrowly defined syllabic rate.

Finally, although the speech stimuli were synthesized to be as natural as possible and were even presented over VR and through real speakers, they were not natural sounding. There were no prosodic cues in these stimuli, such as pitch fluctuations (other than the target events which comprised the task) or changes in emphasis. There was no syntactic or semantic information carried in the vowel streams. The viewing distance was 2 m, which exceeds the comfortable conversational distance in many cases (but was chosen to match the distance of the loudspeaker). Finally, artificial application of vowel formants created minor signal artifacts. These factors combined lead our stimuli to have an effect that observers most commonly described as “creepy”.

### Future directions

In every study of audiovisual speech perception, choices in stimulus design necessarily impact how behavioral results are interpreted. In our previous paper regarding this topic, we separated the relative contributions of temporal coherence and articulatory correspondence by simply rotating the faces, which offered us a good deal of naturalism but very little control over the stimuli (Fiscella et al., 2022). Here, we sacrificed naturalism for control. Development of stimuli that lay in a more balanced place of naturalism and control, in combination with the present work, could help us resolve our understanding of the cues that aid binding in speech.

Our speech stimuli only included two talkers – one male, and one female. Although we find a strong effect of talker identity, it is unclear whether this is due simply to the differing genders of the talker or to the individuals themselves. Additional work, in which multiple talkers have both similar and different vocal characteristics and gender presentation, will not only help resolve the question of whether gender is primarily responsible for the performance difference, but also allow us to again look at potential influence of temporal coherence when talkers from the same gender are not so easily segregated.

## Conclusion

Audiovisual binding has the potential to improve perception of speech when we see the face of the talker we are attending. Here, we aimed to understand which audiovisual correspondences of speech might support audiovisual binding, focusing on temporal coherence, linguistic congruence, and semantic congruence. Despite previous studies consistently showing a relationship between temporal coherence and orthogonal event detection, evidence of audiovisual binding, we found no effect of temporal coherence in our task. Instead, we found a strong effect of the identity of the face and voice, which reduced false alarms when matched.

Our results call into question two assumptions made in our task design: 1) binding is the only mechanism by which a participant can improve their performance in this task, 2) the attentional demands of performing a dual task with a single masker will place the subject in a task-difficulty regime where audiovisual binding is helpful. We now suggest that seeing the face of the talker that matches the target auditory stream acts as a high-level cue to aid in selecting the correct voice. The present results emphasize the need to better understand the neural underpinnings of effects that we have labeled as “audiovisual binding” as well as to refine the tasks we use to study it, ensuring they are truly specific.

## Data Availability

All data and analysis code are available at https://github.com/maddoxlab/vowels_cappelloni_2022.

## Acknowledgements

The authors would like to thank Adrian KC Lee, Jennifer K Bizley, Justin T Fleming, Abigail K Noyce, Liesbeth Gijbels, and other attendees for their active participation in a discussion which guided the framing of our results.

